# Two distinct scene processing networks connecting vision and memory

**DOI:** 10.1101/057406

**Authors:** Christopher Baldassano, Andre Esteva, Diane M. Beck, Li Fei-Fei

## Abstract

A number of regions in the human brain are known to be involved in processing natural scenes, but the field has lacked a unifying framework for understanding how these different regions are organized and interact. We provide evidence from functional connectivity and meta-analyses for a new organizational principle, in which scene processing relies on two distinct networks that split the classically defined Parahippocampal Place Area (PPA). The first network consists of the Occipital Place Area (OPA/TOS) and posterior PPA, which contain retinotopic maps and are related primarily to visual features. The second network consists of the caudal Inferior Parietal Lobule (cIPL), Retrosplenial Cortex (RSC), and anterior PPA, which connect to the hippocampus and are involved in a much broader set of tasks involving episodic memory and navigation. This new framework for understandingthe neural substrates of scene processing bridges results from many lines of research, and makes specific functional predictions.

## Introduction

Natural scene perception has been shown to rely on a distributed set ofcortical regions, including the parahippocampal place area (PPA) (Epstein & Kanwisher, 1998), retrosplenial cortex (RSC) (O'Craven & Kanwisher, 2000), and the occipital place area (OPA, aka transverse occipital sulcus, TOS) (Hasson, Harel, Levy, & Malach, 2003; Nakamura et al., 2000). More recent work has suggested that the picture is even more complicated, with multiple subdivisions within PPA and the possible involvement of the parietal lobe (Baldassano, Beck, & Fei-Fei, 2013).Although there has been substantial progress in understanding the functional properties of each of these regions and the differences between them, the field has lacked a coherent framework for summarizing the overall architecture of the human scene processing system.

There is a long history of proposals for partitioning the visual systeminto separable components with different functions, such as spatial frequency channels (Campbell & Robson, 1968), what versus where/how pathways (Kravitz, Saleem, Baker, & Mishkin, 2011; Mishkin, Ungerleider, & Macko, 1983), or magnocellular, parvocellular, and koniocellularstreams (Kaplan, 2004). With respect to natural scene perception, one can imagine at least two separable functions: processing the specific visual features present in the current glance of a scene, and connecting that to the stable, high-level knowledge of where the place exists in the world, what has happened here in the past, and what possible actions we could take here in the future. For most cognitive and physical tasks we undertake in real-world places, the specific visual attributes we perceive are just a means to this end, of recalling and updating information about the physical environment; “the essential feature of a landmark is not its design, but the place it holds in a city's memory” (Muschamp, 2006). The connection between place and memory has been recognized for thousands of years, reflected in the ancient Greek “method of loci”that strengthens a memory sequence by associating it with physical locations (Yates, 1966).

Some previous work has begun to point to this type of organizing principle among scene perception regions. Mapping functional connectivity differences between pairs of scene-sensitive regions has revealed some consistent distinctions, with some regions more connected to visual cortex and others connected to parietal and medial temporal regions (Baldassano et al., 2013; Nasr, Devaney, & Tootell, 2013). Contrasting activity evoked by perceptual categorization tasks compared to semantic retrieval tasks shows a similar division between visual and higher-level cortex (Fairhall, Anzellotti, Ubaldi, & Caramazza, 2014). These experiments, however, have all been targeted, hypothesis-driven comparisons between regions with similar functional properties. It is unclear whether these divisions are major organizing principles of the brain’s connectivity networks, orsimply subtle differences within asingle coherent scene-processing network.

To answer this question, we took a data-driven approach to identifying scene-sensitive regions and clustering cortical connectivity. We first aggregate local high-resolution resting-state connectivity information into spatially-coherent parcels (Baldassano, Beck, & Fei-Fei, 2015), in order to increase signal to noise and obtain more interpretable units than individual voxels. We then apply hierarchical clustering to show that there exists a natural division in posterior human cortex that splits scene-related regions into two separate, bilaterally-symmetric networks. The posterior network includes OPA and the posterior portion of PPA (retinotopic maps PHC1 and PHC2), while the anterior network is composed of the RSC, anterior PPA (aPPA), and the caudal inferior parietal lobule (cIPL). We then show that these two networks differ in their connectivity to the hippocampus, with the anterior network exhibiting much higher resting-state hippocampal coupling (especially to anterior hippocampus), suggesting that memory-and navigation-related functions are primarily restricted to the anterior network. We provide supporting evidence for this functional division from a reverse-inference meta-analysis of previous results from visual, memory,and navigation studies, and an atlas of retinotopic maps.

Based on these results, as well as a review of previous work, we propose that scene processing is fundamentally divided into two collaborating but distinct networks, with one focused on the visual features of a scene image and the other related to contextual retrieval and navigation. Under this framework, scene perception is less the function of a unified set of distributed neural machinery and more of “an ongoing dialogue between the material and symbolic aspects of the past and the continuously unfolding present” (Baker, 2012).

## Results

Our primary dataset is a 1.8-billion element resting-state connectivity matrix distributed by the Human Connectome Project (Van Essen et al., 2013), which estimates the timecourse correlation between every pair of locations in the brain at 2mm resolution based on a group of 468 subjects. Since we wish to understand the large-scale structure of visual cortex, it is helpful to abstract away from individual voxels and study the functional and connectivity properties of larger parcels. Rather than imposing a parcelltion based on specific regions of interest, we used a data-driven approach to produce spatially-coherent parcels tiling the cortical surface in a way that retains as much information as possible from the full connectivity matrix (Baldassano et al., 2015). This parcellation consists of 172 regions across both hemispheres, each of which contains surface vertices that all have very similar connectivity patterns with the rest of the brain. The connectivity matrix between these 172 parcels captures more than 76%of the variance in the original connectivity matrix, despite being dramatically smaller (by five orders of magnitude). A visualization of this connectivity space is shown in Supplementary Figure 1.

### Clustering Parcels into Networks

To determine how these local parcels are organized into distributed networks, we performed hierarchical clustering to group together parcels with high functional connectivity (regardless of their spatial position). These networks are remarkably similar between hemispheres (despite not being constrained to be symmetric), as shown in the 10-network clustering in Figure 1.

Which of these networks are directly related to scene perception? We used data from a standard localizer in a separate group of subjects to define group-level regions of interest for scene-selective regions OPA, PPA, and RSC. We also anatomically identified cIPL as in previous work, since this region has been shown to have functional connections to scene regions (Baldassano et al., 2013).

We found that these scene ROIs fell almost entirely onto two of the connectivity networks. A posterior network (dark blue), overlapping OPA and posterior PPA, covered all of visual cortex outside of an early foveal cluster. An anterior network (magenta), overlapping cIPL, RSC, and anterior PPA, covered a parietal/medial-temporal network that includes anterior temporal and orbitofrontal parcels. This corresponds to a portion of the known default mode regions, with other default mode regions being grouped into aseparate network (green); a similar fractionation of the default mode has been proposed previously (Andrews-Hanna, Reidler, Sepulcre, Poulin, & Buckner, 2010).

We note that this bifurcation into posterior and anterior scene networks occurs between neighboring parcels in both dorsal and ventral regions; i.e. there is a rapid change in connectivity profiles between OPA and cIPL and along the posterior-anterior axis of PPA. To statistically evaluate (in individual subjects) the shift in network membership in the dorsal parcels near OPA/cIPL, we measured functional connectivity between these parcels and a reference parcel in the anterior network. According to our two-network hypothesis, we predict increasing connectivity to this anterior-network reference parcel as we move from posterior to anterior parcels within both the dorsal and ventral stream. In order to avoid influences from local noise correlations, we selected the reference parcel to be the parcel near RSC on the medial surface. Similarly, we measured connectivity between the ventral parcels near PPA and a reference parcel on the opposite lateral surface (the most anterior dorsal parcel, overlapping cIPL).

In both cases, we observed rapid increases in connectivity as we moved posterior to anterior across the network boundaries (Figure 2). Along the dorsal boundary, we see significant increases in connectivity to the RSC parcel when moving from the first to the second parcel (Left: t_19_=6.98, p<0.001; Right: t_19_=6.35, p<0.001; two-tailed paired t-test), from the second to the third parcel (Left: t_19_=7.72, p<0.001; Right: t_19_=6.16, p<0.001), and from the third to the fourth parcel (Right: t_19_=2.44, p=0.025). We observe a similar significant (though less dramatic) increase in connectivity to the cIPL parcel when moving from the first to the second PPA parcel (Left: t_19_=4.21, p<0.001; Right: t_19_=2.68, p=0.015) and from the second to the third PPA parcel (Right: t_19_=3.03, p=0.007).

**Figure 1.**
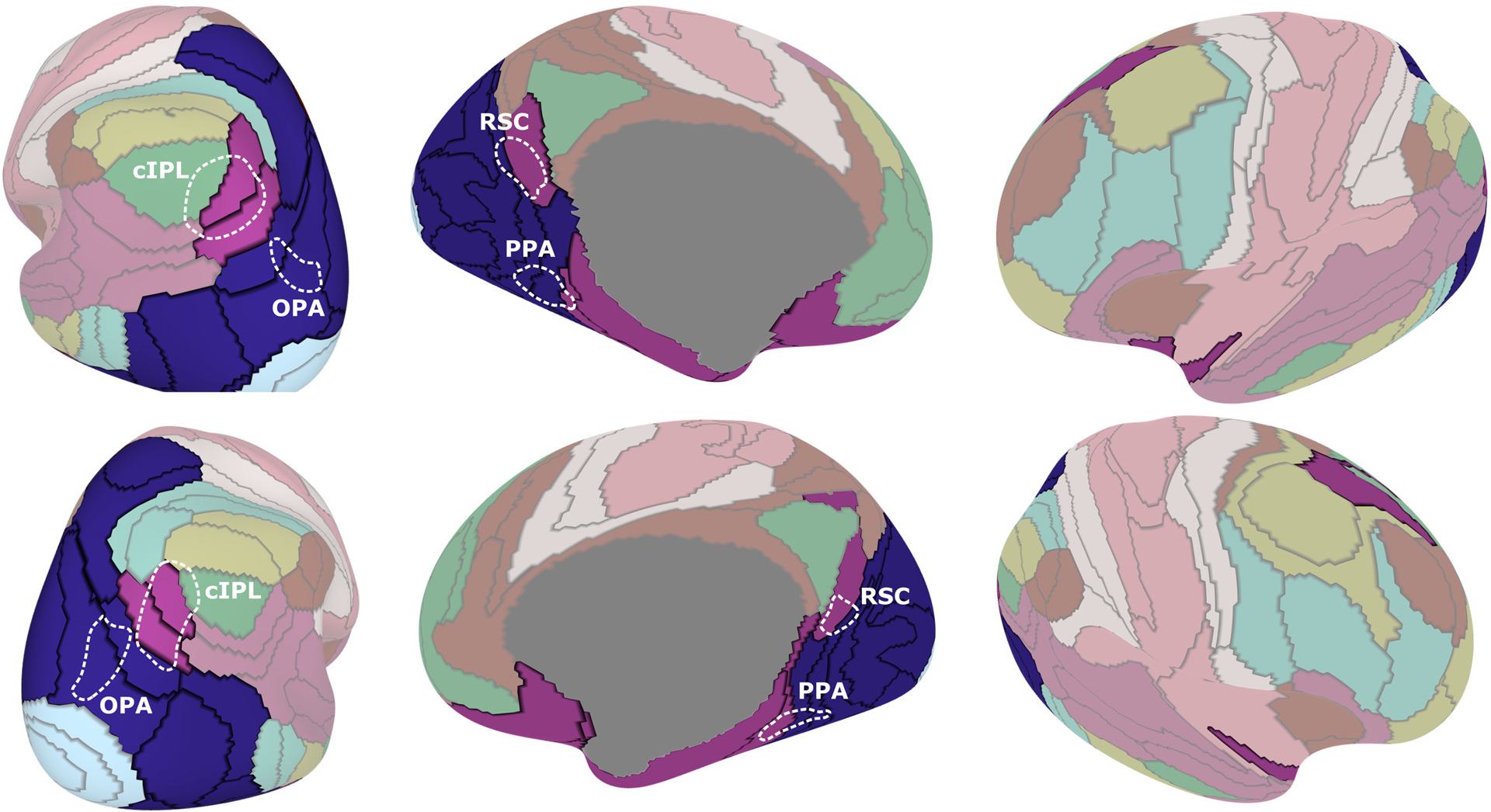
Connectivity clustering of cortical parcels. The cortex was first grouped into 172 local parcels (black lines) such that the surface vertices in each parcel had similar connectivity properties. Performing a second-level hierarchical clustering to group together strongly-connected parcels shows that scene-related regions are split across two networks, which are largely symmetric across left (top row) and right (bottom row) hemispheres. OPA and posterior PPA overlap with a posterior network(dark blue) that covers all of visual cortex outside the foveal confluence, while cIPL, RSC, and anterior PPA overlap with an anterior network (magenta) that covers much of the default mode network.

**Figure 2.**
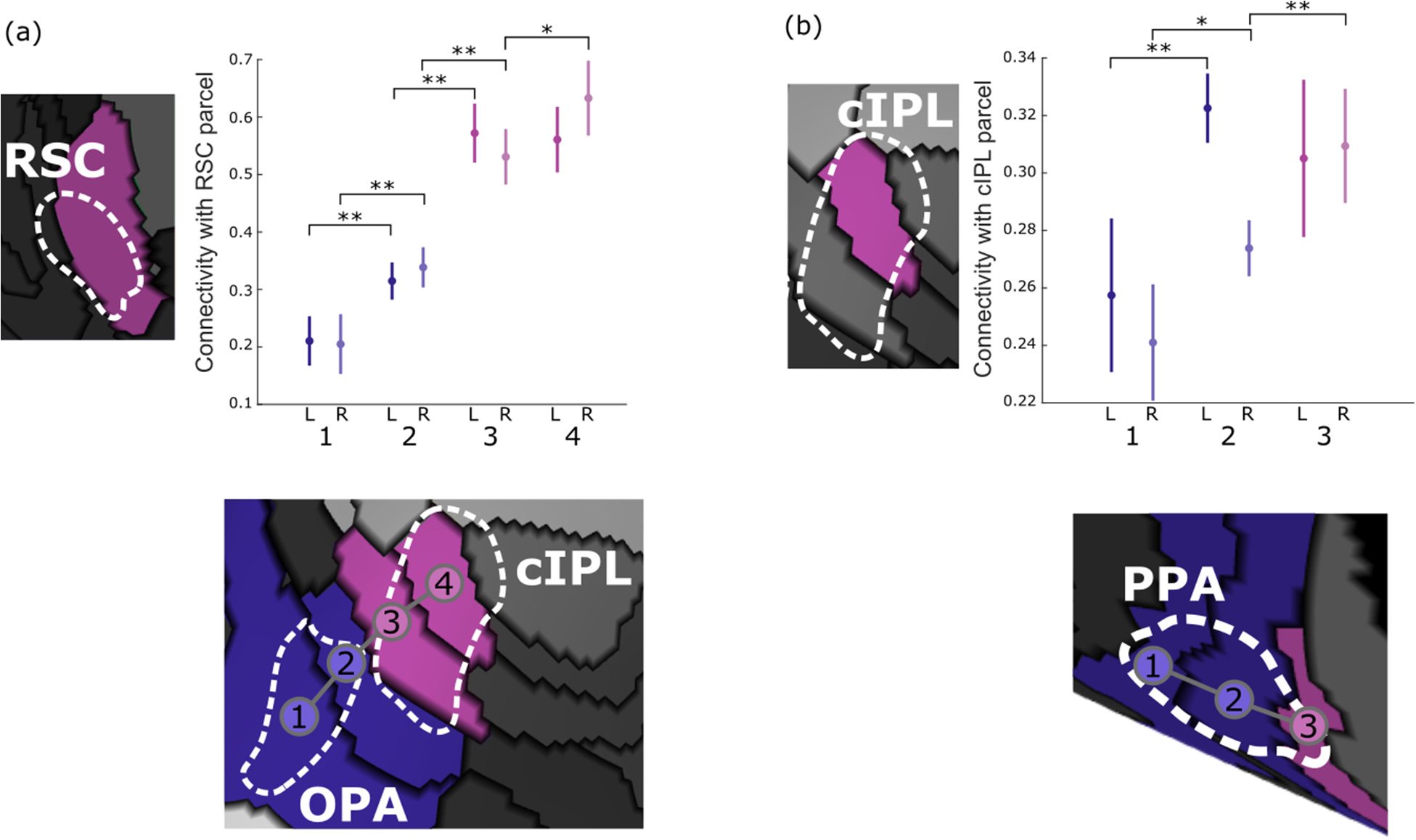
Connectivity shifts across the network border. (a) Dorsal parcels markedly increase their connectivity to the medial RSC parcel as we move anteriorly from parcels overlapping OPA to parcels overlapping cIPL, indicating a sharp transition between networks near the OPA/cIPL border.(b) Ventral parcels also show a shift in network connectivity properties, with increasing connectivity to the anterior cIPL parcel as we move from pPPA to aPPA. Error bars are 95% confidence intervals across subjects, *p<0.05,**p<0.01.

### Connectivity with the Hippocampus

Since the anterior scene-network overlaps with default mode regions, while the posterior scene-network does not, we predict that the anterior network should be more related to memory and navigation tasks that engage the hippocampus. To test this hypothesis, we measured the functional correlation at rest between mean hippocampal activity and the mean activity in each parcel within the posterior and anterior scene networks. As shown in Fig. 3, there is a dramatic difference in hippocampal connectivity for parcels in the posterior network (overlapping with OPA and posterior PPA) compared to the anterior network (overlapping with RSC, cIPL, and anterior PPA).Moving posterior to anterior along the dorsal path, hippocampal connectivity first decreases slightly (first parcel to second parcel, Left: t_19_=-3.04, p=0.007; Right: t_19_=2.15 p<0.04, two-tailed paired t-test), then increases significantly when moving to the third parcel (Left: t_19_=5.62, p<0.001; Right: t_19_=3.79, p=0.001) and to the fourth parcel (Left t_19_=4.17, p<0.001; Right: t_19_=5.74, p<0.001). Along the ventral path, hippocampal connectivity jumps from the first to the second parcel overlapping with PPA (Left: t_19_=5.27, p<0.001; Right: t_19_=5.76, p<0.001) and from the second to the third parcel (Right: t_19_=5.80, p<0.001). This elevated hippocampal connectivity in the anterior network parcels is consistent with our hypothesis that the anterior network is more closely related to navigation and memory.

We also investigated whether this effect was being driven by a subregion of the hippocampus, by correlating the mean timecourse in both scene networks with the timecourses of each posterior-to-anterior coronal slice of the hippocampus. Our results show that the entire hippocampus is more strongly connected to the anterior scene-network than the posterior scene-network, but this difference is especially large in the anterior hippocampus. To confirm this pattern of results, we divided the hippocampus into posterior and anterior subregions at y=-21 (Zeidman, Mullally, & Maguire, 2015) and correlated their mean timecourses with the two scene-network timecourses. This analysis confirmed that the anterior network is more strongly connected to both posterior (t_19_=7.66, p<0.001, two-tailed paired t-test) and anterior (t_19_=6.58, p<0.001) hippocampus, and that this anterior-network connectivity is larger in anterior hippocampus (t_19_=3.29, p=0.004); a repeated-measures ANOVAshows significant main effects of bothhippocampal subregion (F_1,19_=11.32, p=0.003) and scene network (F_1,19_=59.2, p<0.001), and an interaction (F_1,19_=7.03, p=0.016). Note that both the anterior and posterior scene networks are closer to posterior hippocampus,ruling out a distance-based explanation for this pattern of results.

**Figure 3.**
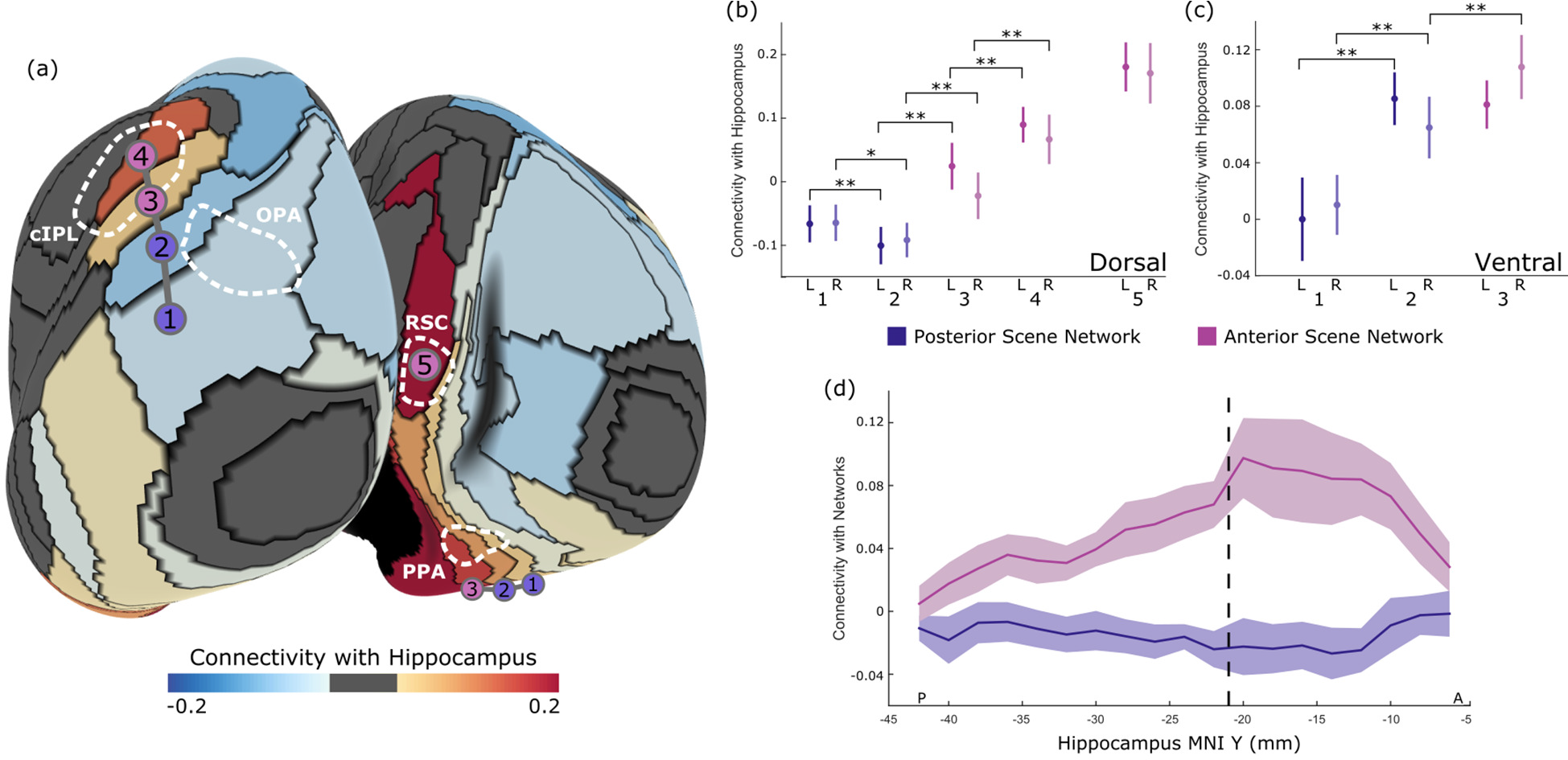
Connectivity between network parcels and the hippocampus. (a) For each parcel in the anterior and posterior scene networks, we computed its resting-state connectivity with the hippocampus, showinga striking increase in hippocampal activity for anterior network parcels overlapping with cIPL, RSC, and anterior PPA (magenta circles) compared to posterior network parcels (blue circles). (b) Along the dorsal network boundary, hippocampal activity first dips slightly and then increases substantially, becoming strongest in the most anterior parcel intersecting cIPL (and is also high in RSC). (c) Ventrally along parcels overlapping with PPA, we observe a similar posterior-anterior gradient in connectivity. (d) Computing the connectivity between each coronal slice of the hippocampus andthe twoscene networks shows that this increased coupling to the anterior network is present throughout the hippocampus, but is especially pronounced in anterior hippocampus (MNI y>-21mm). Error bars are 95% confidence intervals across subjects, *p<0.05,**p<0.01.

### Comparison to Meta-analyses and Retinotopic Atlas

The connectivity results described thus far suggest a functional division for scene-related regions, with some belonging to a posterior network and others belonging to an anterior network. To assess the functional significance of these two networks, we ran two reverse-inference meta-analyses using the-*NeuroSynth* tool (Yarkoni, Poldrack, Nichols, Van Essen, & Wager, 2011). This system automatically extracts activation coordinates from many fMRI studies (greater than 10,000 at the time of writing); given a particular set of studies, it can identify voxels that are*more likely* to beactivated in this set of studies *relative to the full set of studies.* These voxels are therefore preferentially active in the query set compared to general fMRI experiments. Based on the areas involved, we hypothesize that the posterior network processes the current visual properties of the scene, whereas the anterior network incorporates episodic memories and contextual aspects of the scene. Thus, in Fig. 4a, we compare meta-analyses for the query “scene” (47 studies) with the query “episodic memory OR navigation OR past future” (125 studies). Along the parahippocampal gyrus,we find that the visual scene activations tend to be posterior to the memory activations, and that the transition point corresponds almost exactly to the division between our two networks. Dorsally, we also observe a separation between the reverse inference maps, with scene and memory activations falling into our two separate networks. Overall, voxels significant only in the scene meta-analysis were concentrated in the posterior network (66% in posterior network, 18% in anterior, 16% in other)while voxels significant only inthe memory/navigation meta-analysis were spread more widely across the cortex, but were concentrated more in the anterior than posterior network (16% posterior, 42% anterior, 42% other). Voxels significant in both the scene and memory/navigation meta-analyses tended to fall near the border between the two networks and divided roughly equally between them (44% posterior, 53% anterior, 4% other).

Another prediction of our framework is that voxels whose activity is tied to specific locations in the visual field (i.e. retinotopic) should, as clearly visual voxels, be present only in the posterior scene network. In Figure 4b, we compared our networks to a group-level probabilistic atlas of retinotopic visual field maps (Wang, Mruczek, Arcaro, & Kastner, 2014). The vast majority of the probability mass in this atlas is concentrated in the posterior network. In early visual cortex (V1, V2, V3, hV4)all non-foveal portions of the visual field maps fall in the posterior network (80% posterior, 0% anterior, 20% other). Ventrally, the posterior network covers VO1/2 (100% posterior, 0% anterior, 0% other), PHC1 (98% posterior, 2% anterior, 0% other), and the peak of the probability distribution for PHC2, which also extends slightly across the anterior network border (78% posterior, 22% anterior, 0% other). Laterally and dorsally, the posterior network includes most ofthe LO1/2 and TO1/2 maps (82%posterior, 0% anterior, 17% other), V3a and V3b (96% posterior, 0% anterior, 3% other), and IPS0-IPS5 (68%posterior, 4% anterior, 28% other), with SPL1 being the only map falling substantially outside the networks we consider (18% posterior, 2% anterior, 80% other).

**Figure 4.**
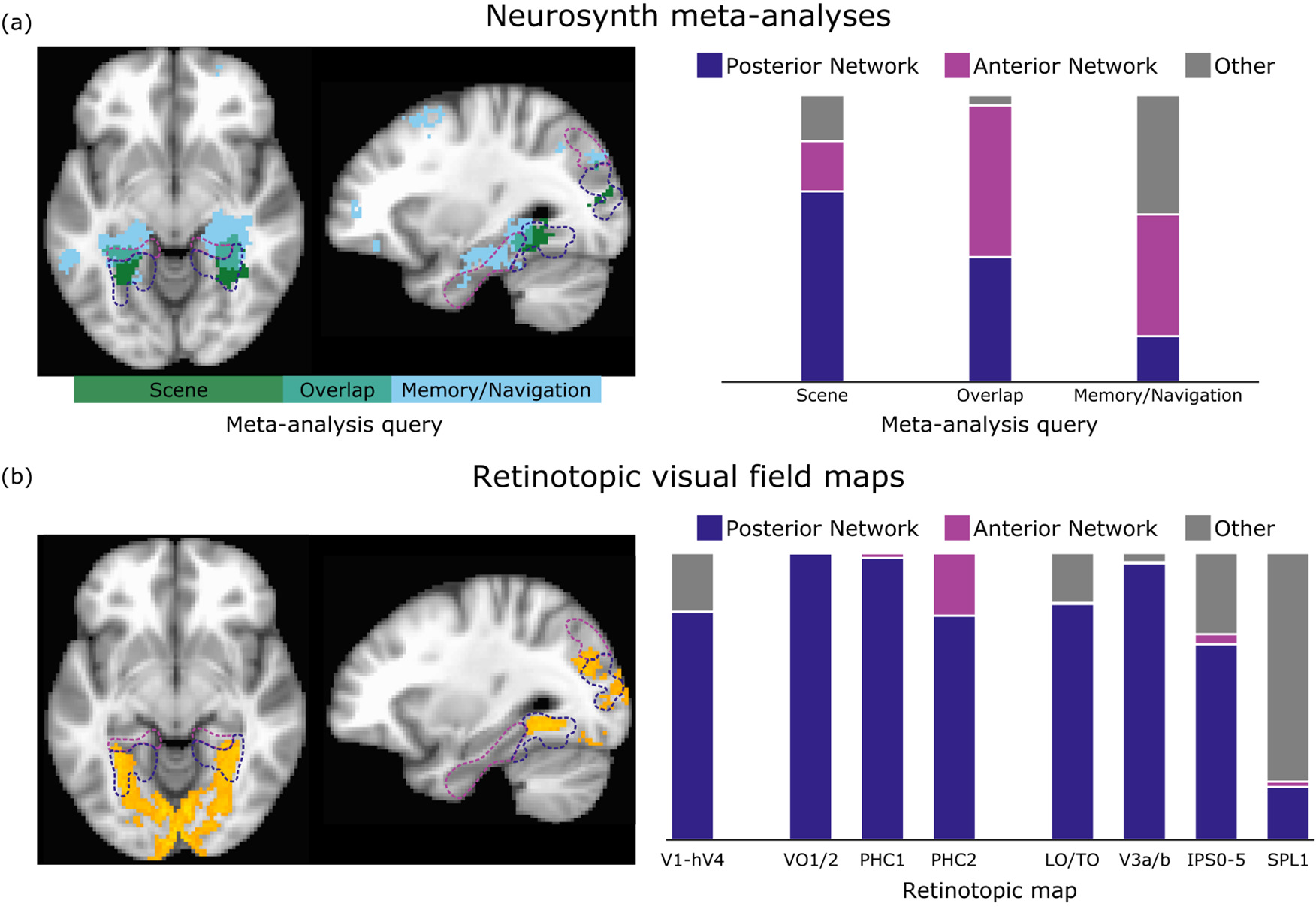
Overlap of posterior and anterior scene networks with previous work. (a)Two meta-analyses conducted using *NeuroSynth* identified overlapping but distinct reverse-inference maps corresponding to studies of visual scenes and to studies of higher-level memory and navigation tasks. These maps separate into our two scene networks, with visual scenes activating voxels in the posterior network and memory/navigation tasks activating voxels in the anterior network, as shown on example axial (z=-8) and sagittal (x=-30) slices. FDR<0.01, cluster size 80 voxels (640 mm^3^). (b) Voxels having a greater than 50%chance of belong to a retinotopic map (orange) overlap with much of the posterior scene network, but end near the border of the anterior scene network. Breaking up the contributions of individual regions, we find that the probability mass of the topographic maps falls primarily within the posterior network, with only PHC2 showing a small overlap with the anterior network (probabilistically at the group level).

### Discussion

**Figure 5.**
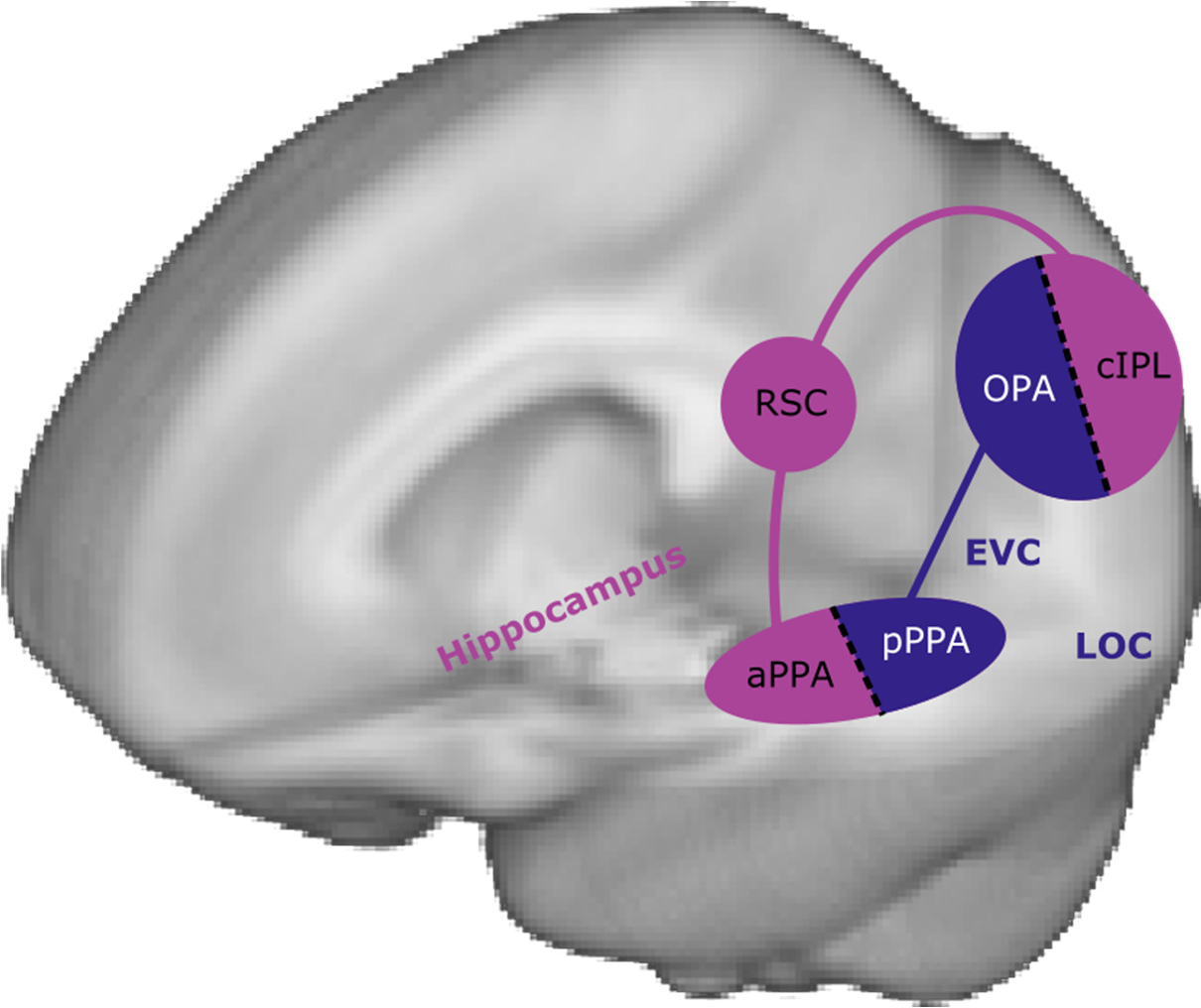
Two-network model of scene perception. Our results provide strong evidence for dividing scene-sensitive regions into two separate networks. We argue that OPA and posterior PPA (PHC1/2) process the current visual features of a scene (in concert with other visual areas, such early visual cortex, EVC, and lateral occipital cortex, LOC), while cIPL, RSC, and anterior PPA perform higher-level context and navigation tasks (drawingon long-term memory structures such as the hippocampus).

By combining a variety of data sources, we have shown converging evidence for a functional division of scene-processing regions into two separate networks (summarized in Figure 5). The posterior visual network covers retintopically-organized regions including OPA and posterior PPA (pPPA), while an anterior memory-related network connects cIPL, RSC, and anterior PPA (aPPA). This division emerges from a purely data-driven network clustering, suggesting that this is a core organizing principle of the visual system.

### Subdivisions of the PPA

The division of the PPA into multiple anterior-posterior subregions with differing connectivity properties replicates our previous work (Baldassano et al., 2013) on an entirely different large-scale dataset, and shows that there is a strong connection between connectivity changes in PPA and the boundaries of retinotopic field maps. There is now a growing literature on anterior versus posterior PPA, including not only connectivity differences (Nasr et al., 2013) but also the response to low-level (Baldassano, Beck, & Fei-Fei, 2016; Baldassano, Fei-Fei, & Beck, 2016; Nasr, Echavarria, & Tootell, 2014; Silson E. H., Chan, Reynolds, Kravitz, & Baker, 2015; Watson, Hymers, Hartley, & Andrews, 2016) and high-level (Aminoff & Tarr, 2015; Linsley & Macevoy, 2014; Marchette, Vass, Ryan, & Epstein, 2015; Park, Konkle, & Oliva, 2014) scene features, as well as stimulation studies (Rafique, Solomon-Harris, & Steeves, 2015). Our results place this division into alarger context, and demonstrate that the connectivity differences within PPA are not just an isolated property of this region buta general organizing principle for scene-processing regions.

### The visual network

The visual network shows a close correspondence with the full set of retinotopic maps identified in previous studies (Brewer & Barton, 2012; Huang & Sereno, 2013; Wang et al., 2014). Previous measurements in individual subjects have also shown strong overlap between OPA and retinotopic maps, especially V3b and LO2 (Bettencourt & Xu, 2013; Nasr et al., 2011; Silson E. H., Groen, Kravitz, & Baker, 2016), and between pPPA and VO2, PHC1, and PHC2 (Arcaro, McMains, Singer, & Kastner, 2009). The only portion of cortex with known retinotopic maps that is not clustered in this network is the shared foveal representation of early visual areas, which segregates into its own cluster, consistent with other work showing a peripheral eccentricity bias in the scene network (Baldassano, Fei-Fei, et al., 2016; Goesaert & Op de Beeck, 2010; Huang & Sereno, 2013; Malach, Levy, & Hasson, 2002).

OPA and posterior PPA have been shown to be closely related to the visual content of a stimulus. Even low-level manipulations of spatial frequency (Kauffmann, Ramanoël, Guyader, Chauvin, & Peyrin, 2015; Rajimehr, Devaney, Bilenko, Young, & Tootell, 2011; Watson et al., 2016) or rectilinearity (Nasr et al., 2014) can drive responses in these regions. Higher-level visual features also drive response patterns in these regions (Bryan, Julian, & Epstein, 2016), and they are hypothesized to be involved in extracting visual environmental features that can be used for navigation (Julian, Ryan, Hamilton, & Epstein, 2016; Marchette et al., 2015). However, neither OPA nor posterior PPA show reliable familiarity effects (Epstein et al., 2007; see further discussion below).

The functional distinction between pPPA and OPA is currently unclear. Previous work has speculated about the purpose of the apparent ventral and dorsal “duplication” of regions sensitive to large landmarks, proposing that it may be related to different output goals (e.g. action planning in OPA, object recognition in pPPA) (Konkle & Caramazza, 2013), or to different input connections (e.g. lower visual field processing in OPA, upper visual field processing in pPPA) (Kravitz, Saleem, Baker, Ungerleider, & Mishkin, 2013; Silson E.H. et al., 2015). OPA and pPPA may also use information from different visual eccentricities, with OPA processing less peripheral, relatively high-resolution environmental features and pPPA processing more peripheral, large-scale geometry and context (Baldassano, Fei-Fei, et al., 2016).

### The memory and navigation network

The network of parahippocampal, retrosplenial, and posterior parietal regions we identify has been emerged independently in many different fields of neuroimaging, outside of scene perception. Meta-analyses of internally-directed tasks such as theory of mind, autobiographical memory, and prospection have identified this as a core, re-occurring network (Kim, 2010; Spreng, Mar, & Kim, 2009; component C10 of Yeo et al., 2014). It comprises a subset of the broader default mode regions, but functional and anatomical evidence suggests that it is a distinct, coherent subnetwork (Andrews-Hanna et al., 2010; Andrews-Hanna, Smallwood, & Spreng, 2014; Yeo et al., 2011). The broad set of tasks which recruit this network have been summarized in various ways, such as “scene construction” (Hassabis & Maguire, 2007), “mnemonic scene construction” (Andrews-Hanna et al., 2010), “long-timescale integration” (Hasson, Chen, & Honey, 2015), or “relational processing” (Eichenbaum & Cohen, 2014). A review of memory studies referred to this network as the posterior medial (PM) memory system, and proposed that it is involved in any task requiring “situation models” relating entities, actions, and outcomes (Ranganath & Ritchey, 2012).

The network has strong functional connections to the hippocampus, which has been implicated in a broad set of cognitive tasks involving “cognitive maps” for organizing declarative memories, spatial routes, and even social dimensions (Eichenbaum & Cohen, 2014; Schiller et al., 2015). During perception, the hippocampus binds together visual elements of an image (Olsen, Moses, Riggs, & Ryan, 2012; Warren, Duff, Jensen, Tranel, & Cohen, 2012; Zeidman et al., 2015), which is especially important for scene stimuli (Graham et al., 2006; Hodgetts, Shine, Lawrence, Downing, & Graham, 2016; Lee, Buckley, et al., 2005; Lee, Bussey, et al., 2005), and then stores this representation into long-term memory (Ryan & Cohen, 2004). As we become familiar with an environment, the hippocampus builds a map of the spatial relationships between visual landmarks, which is critical for navigation (Morgan, Macevoy, Aguirre, & Epstein, 2011). Recalling or even imagining scenes also engages the hippocampus,especially anterior hippocampus, which may serve to integrate memory and spatial information (Zeidman & Maguire, 2016). Our results argue that only the anterior scene regions are directly involved in building hippocampal representations of the environment, and in retrieving relevant memories and navigational information for a presented or imagined scene.

The specific functions of the individual components of this network have also been studied in a number of contexts. RSC appears to be most directly involved in orienting the viewer to the structure of the environment (both within and beyond the borders of the presented image) for the purpose of navigational planning; it encodes both absolute location and facing direction (Epstein & Vass, 2014; Marchette, Vass, Ryan, & Epstein, 2014; Vass & Epstein, 2013), integrates across views presented in a panoramic sequence (Park & Chun, 2009), and shows strong familiarity effects (Epstein et al., 2007; Epstein, Parker, & Feiler, 2007). This is consistent with rodent neurophysiological studies, which have identified head direction cells in this region (Chen, Lin, Green, Barnes, & McNaughton, 1994). RSC is not sensitive to low-level rectilinear features in non-scene images such as objects or textures (Nasr et al., 2014), though it does show some preference for rectilinear features in images of 3D scenes (Nasr et al., 2014; Watson et al., 2016).

The specific properties of anterior PPA have been less well-studied, since it was not recognized as a separate region within the PPA until recently. It has been shown to be driven more by high-level category information than spatial frequency content (Watson et al., 2016), to represent real-world locations (even from perceptually-distinct views) (Marchette et al., 2015), to encode object co-occurrences (Aminoff & Tarr, 2015), and to represent real-world physical scene size (Park et al., 2014). Its representation of scene spaciousness draws on prior knowledge about the typical size of different scene categories, since it is affected by the presence of diagnostic objects (Linsley & Macevoy, 2014).

The cIPL (also referred to as pIPL, PGp, or the angular gyrus) has been proposed as a “cross-modal hub” (Andrews-Hanna et al., 2014)that connects visual information with other sensory modalities as well as knowledge of the past. It is more intimately associated with visual cortex than most lateral parietal regions, since it has strong anatomical connections to higher-level visual regions in humans and macaques (Caspers et al., 2011), and has a neurotransmitter receptor distribution similar to V3v and distinct from the rest of the IPL (Caspers et al., 2012). It has been mostly ignored in the scene perception literature, primarily because it is not strongly responsive to standard scene localizers that show sequences of unfamiliar and unrelated scene images. For example, a study showing familiarity effects in cIPL described this location only as “near TOS” (Epstein et al., 2007). The cIPL appears commonly, however, in studies involving personally familiar places, which are associated with a wealth of memory, context, and navigational information. It is involved in memory for visual scene images (Elman, Rosner, Cohn-Sheehy, Cerreta, & Shimamura, 2013; Montaldi, Spencer, Roberts, & Mayes, 2006; Takashima et al., 2006; van Assche, Kebets, Vuilleumier, & Assal, 2014), learning navigational routes (Bray, Arnold, Levy, & Iaria, 2014; Burgess, Maguire, Spiers, & O'Keefe, 2001), and even imagining past events or future events in familiar places (Hassabis, Kumaran, & Maguire, 2007; Szpunar, Chan, & McDermott, 2009). It can integrate information across space (Livne & Bar, 2016) and time (Lerner, Honey, Silbert, & Hasson, 2011; Vilberg & Rugg, 2012), and has been shown in lesion studies to be critical for orientation and navigation (Kravitz et al., 2011). Our connectivity results and meta-analysis suggest that cIPL may play a prominent role in connecting visual scenes to the real-world location they depict.

### Contrasting the two networks

Although our work is the first to propose the visual versus context networks as a general framework for scene perception, several previous studies have shown differential effects within these two networks. Contrasting the functional connectivity patterns of RSC vs. OPA or LOC (Nasr et al., 2013) or anterior vs. posterior PPA (Baldassano et al., 2013) show a division between the two networks, consistent with our results. Contrasting scene-specific activity with general (image or word) memory retrieval showedananterior vs. posterior distinction in PPA and cIPL/OPA, with only more anterior regions (aPPA and cIPL, along with RSC) responding to content-independent retrieval tasks (Fairhall et al., 2014; Johnson & Rugg, 2007). Our two-network division is also consistent with the “dual intertwined rings” model, which argues for a high-level division of cortex into a sensory ring and an association ring, the second of which is distributed but connected into a continuous ring through fiber tracts (Mesmoudi et al., 2013).

### Open questions

The anterior/posterior pairing of aPPA/pPPA and cIPL/OPA raises the question of whether there is a similar anterior/posterior division in RSC. There is some evidence to suggest that this is the case: wide-field retinotopic mapping using natural scenes shows a partial retinotopic organization in RSC (Huang & Sereno, 2013), and RSC's response to visual rectilinear features appears to be limited to the posterior portion (Nasr et al., 2014). However a study of retinotopic coding in scene-selective regions failed to find any consistent topographic organization to RSC responses (Ward, MacEvoy, & Epstein, 2010), and previous analyses of the functional properties of anterior versus posterior RSC have not found any significant differences (Park et al., 2014). More precise functional mapping of the regions near the parieto-occipital sulcus will be required to see if a retinotopic subregion is present in this scene-sensitive region as well.

Another interesting question is how spatial reference frames differ between and within the two networks. Given its retinotopic fieldmaps, the visual network presumably represents scene information relative to the current eye position; previous work has argued that this reference frame is truly retina-centered and not egocentric (Golomb & Kanwisher, 2012; Ward et al., 2010). The context network, however, likely transforms information between multiple reference frames. Models of spatial memory suggest that medial temporal lobe (possibly including aPPA) utilizes an allocentric representation, while the posterior parietal lobe (possibly including cIPL)is based on an egocentric reference frame, and that the two are connected via a transformation circuit in RSC that combines allocentric location and head direction (Byrne, Becker, & Burgess, 2007; Vann, Aggleton, & Maguire, 2009). There is some recent evidence for this model in human neuroimaging: posterior parietal cortex codes the direction of attention in an egocentric reference frame (even for positions outside the field of view) (Schindler & Bartels, 2013), and RSC contains both position and head direction information (anchored to the local environment) (Marchette et al., 2014). This raises the possibility that another critical role of cIPL could be to transform retinotopic visual information intoa stable egocentric scene over the course of multiple eye movements. The properties of aPPA, however, are much less clear; it seems unlikely that it would utilize an entirely different coordinate system than neighboring PHC1/2, and some aspects of the scene encoded in aPPA such as object co-occurrence (Aminoff & Tarr, 2015) do not seem tied to any particular coordinate system.

### Conclusion

Based on data-driven connectivity analyses and analysis of previous literature, we have proposed a unifying framework for understanding the neural systems involved in processing both visual and non-visual properties of natural scenes. This new two-network classification system makes explicit the relationships between known scene-sensitive regions, re-emphasizes the importance of the functional subdivision within the PPA, and incorporates posterior parietal cortex as a primary component of the scene-understanding system. Our proposal, that much of the scene-processing network relates more to contextual and navigational information than to specific visual features, suggests that experiments with unfamiliar natural scene images will give only a partial picture of the neural processes evoked in real-world places. Experiencing our visual environment requires a dynamic cooperation between distinct cortical systems, to extract information from the currentview of a scene and then integrate it with our understanding of the world and determine our place in it.

## Materials and Methods

### Imaging Data

The majority of the data used in this study was obtained from the HumanConnectome Project (HCP), which provides detailed documentation on the experimental and acquisition parameters for these datasets (Van Essen et al., 2013). We provide an overview of these datasets below.

The group-level functional connectivity data were derived from the 468-subject group-PCA eigenmaps, distributed with the June 2014 “500 Subjects” HCP data release. Resting-state fMRI data were acquired over four sessions (14 min, 33 seconds each) while subjects fixed on a bright cross-hair on a dark background, using a multiband sequence to achieve a TR of 720ms at 2.0mm isotropic resolution (59412 surface vertices). These timecourses were cleaned using FMRIB's ICA-based Xnoiseifier (FIX) (Salimi-Khorshidi et al., 2014), and then the top 4500 eigenvectors for each voxel were estimated across all subjects using Group-PCA (Smith, Hyvärinen, Varoquaux, Miller, & Beckmann, 2014). This data was used to perform the parcellation and network clustering, and to generate whole-brainmaps (i.e. Figs 1,3a)

For the first 20 subjects within the “500 Subjects” release with complete data (subj ids 101006, 101107, 101309, 102008, 102311, 103111, 104820, 105014, 106521, 107321, 107422, 108121, 108323, 108525, 108828, 109123, 109325, 111413, 113922, 120515), we created individual subject resting-state datasets by demeaning and concatenating their four resting-state sessions. This data was used to statistically measure the robustness of connectivity differences observed in the group-level data (i.e. Figs 2,3b-d). We also obtained these subjects' data from the HCP Working Memory experiment, in which they observed blocks of stimuli consisting of faces, places, tools, or body parts. We collapsed across the two memory tasks being performed by participants (target-detection or 2-back detection).

To identify group-level scene localizers, we used data from a separate set of 24 subjects scanned at Stanford University (see below). Each subject viewed blocks of stimuli from six categories: child faces, adult faces, indoor scenes, outdoor scenes, objects (abstract sculptures with no semantic meaning), and scrambled objects. Functional data were acquired with an in-place resolution of 1.56mm, slice thickness of 3mm (with 1 mm gap), anda TR of 2s; a high-resolution (1mm isotropic) SPGR structural scan was also acquired to allow for transformation to MNI space. Full details of the localizer stimuli and acquisition parameters are given in our previous work(Baldassano et al., 2013).

The cIPL was defined using the Eickhoff-Zilles PGp probabilistic cytoarchitectonic map (Eickhoff et al., 2005) as described in our previous work (Baldassano et al., 2013). The hippocampus was divided into anterior and posterior subregions at MNI y=-21, consistent with previous studies (Zeidman et al., 2015).

### Subjects

Scene localizer data was collected from 24 subjects (6 female, ages 22-32, including one of the authors). Subjects were in good health with no past history of psychiatric or neurological diseases, and with normal or corrected-to-normal vision. The experimental protocol was approved by the Institutional Review Board of Stanford University, and all subjects gave their written informed consent.

### Resting-state Parcellation

We generated a voxel-level functional connectivity matrix by correlating the group-level eigenmaps for every pair of voxels and applying the arctangent function. We parcellated this 59412 by 59412 matrix into contiguous regions, using a generative probabilistic model (Baldassano et al., 2015).This method finds a parcellation of the cortex such that the connectivity properties within each parcel are as uniform as possible, making multiple passes over the dataset to fine-tune the parcel borders. We set the scaling hyperparameter jO=3000 to produce a manageable number of parcels,but our results are similar for a wide range of settings for jO (see Supplementary Fig. 2).

### Scene localizers

To identify PPA, RSC, and OPA, we deconvolved the localizer data from the 24 Stanford subjects using the standard block hemodynamic model in AFNI(Cox, 1996), with faces, scenes, objects, and scrambled objects as regressors. The Scenes>Objects t-statistic was used to define PPA (top 300 voxels near the parahippocampal gyrus), RSC (top 200 voxels near retrosplenial cortex), and OPA (top 200 voxels near the transverse occipital sulcus). The ROI masks were then transformed to MNI space, summed across all subjects, and mapped to the closest vertices on the group cortical surface. The cluster denoting highest overlap between subjects was then manually annotated.

### Parcel-to-parcel and hippocampal functional connectivity

The 468-subject eigenmaps distributed by the HCP are approximately equal to performing a singular value decomposition on the concatenated timecourses of all 468 subjects, and then retaining the right singular values scaled by their eigenvalues (Smith et al., 2014). This allows us to treat these eigenmaps as pseudo-timecourses,since dot products (and thus correlations) between eigenmaps approximate the dot products between the original voxel timecourses. Given a parcellation, we computed the group-level connectivity between a pair of regions by taking the mean over all eigenmaps in each region, then correlating these mean eigenmaps and applying the Fisherz-transform (hyperbolic arctangent).We computed subject-level connectivity in the same way, using the resting-state timecourse for each voxel rather than the eigenmap.

Connectivity between cortical parcels and the hippocampus was computed similarly, using eigenmaps (for group data) or timecourses (for subject data) extracted from the hippocampal volume data distributed by the HCP. In order to focus on hippocampal connectivity differences among parcels, we used the mean gray timecourse regression (MGTR) version of the group data and regressed out the global timecourse from the subject data.

### Multidimensional scaling and Network Clustering

The 172 by 172 parcel functional connectivity matrix was converted intoa distance matrix by subtracting every entry from the maximum entry. Classical multidimensional scaling was applied to the distance matrix, and the first three dimensions were used to assign voxels RGB colors (with each color channel scaled to span the full range of 0 to 255 along each axis) and to plot parcels in a 3D space. We performed the same operation on each subject-level matrix as well, and then aligned each subject's 3D pointcloud to the group pointcloud using a procrustes transform. Hierarchical ward clustering (unconstrained by parcel position) was also applied to the distance matrix to compute a hard clustering into 10 networks.

### Meta-analysis and retinotopic field maps

Two reverse-inference meta-analyses were performed using the *NeuroSynth* website (Yarkoni et al., 2011). *NeuroSynth* is a set of open-source python tools for automatically extracting data from fMRI studies and computing activation likelihood maps, and the website hosts these tools (and associated datasets) for public use. Supplying a keyword query will identify all studies in which that query appears frequently, and then analyzes the activations reported in these queried studies In addition to standard “forward inference” maps giving the probability p(activation|query) that a voxel will be activated in these studies, *NeuroSynth* generates “reverse inference” maps giving the probability p(query|activation) that a voxel activation came specifically from this query set. Voxels appearing the reverse inference map therefore appear more often in the query set relative to the full set of (>10,000) fMRI studies in the database. This accounts for base rate differences in how often activation is observed in different brain regions. Our metaanalyses can be viewed online at http://neurosynth.org/analyses/custom/dda0e003-efd0-4cfa/ and http://neurosynth.org/analyses/custom/9e6df59d-02df-4357/.

A volumetric group-level probabilistic atlas (Wang et al., 2014) was used to define retinotopic field maps. We computed the total probability mass of each map that fell within one of our two networks or in other regions of the cortex, and then normalized the sum of the three values to 100%. For visualization, the probability that a voxel belongs to any field map was computed as 1-Π_*i*_(1-*P_i_*) where *p_i_* is the probability that the voxel falls within field map i.

### Parcel scene selectivity

In order to validate that our group-level functional scene localizer (from the Stanford subjects) properly identified scene-related parcels in the HCP subjects, we used data from Working Memory experiment performed by the 20 HCP subjects. We first used a hemodynamic model to associate timepoints within the working memory experiment with specific stimulus categories. We labeled timepoints as corresponding to bodies, faces, places, or tools by constructing a boxcar timecourse denoting when each stimulus category was being displayed, convolving these indicators with the standard SPM hemodynamic response function provided with AFNI (Cox, 1996), rescaling the maximum value to 1, then re-thresholding to a binary indicator. Effectively, this produced a shift of the stimulus blocks by 5.55s to account for hemodynamic delay. The fMRI timecourses were cleaned by regressing out movement (6 degree-of-freedom translation/rotation and derivatives) and constant, linear, and quadratic trends from each run, then normalizing each voxel to have unit variance. Voxel timecourses were then averaged within each parcel, yielding a vector of average parcel activities for each timepoint. The mean parcel activation to scenes was compared to the mean parcel activation to all other categories, and this difference was statistically tested across the 20 subjects for each parcel. The significance threshold was corrected for multiple comparisons using the same false discovery rate (FDR) procedure implemented in AFNI's3dFDR (Cox, 1996). The results are shown in Supplementary Figure 3.

## Acknowledgements

Funding was provided by a National Science Foundation Graduate Research Fellowship (to CB) under grant number DGE-0645962, and Office of Naval Research Multidisciplinary University Research Initiative (to DMB and LF) grant number N000141410671. Data were provided in part by the Human Connectome Project, WU-Minn Consortium (Principal Investigators: David Van Essen and Kamil Ugurbil; 1U54MH091657) funded by the 16 NIH Institutes and Centers that support the NIH Blueprint for Neuroscience Research; and by the McDonnell Center for Systems Neuroscience at Washington University. We thank the Richard M. Lucas Center for Imaging, the Center for Cognitive and Neurobiological Imaging, and Michael Arcaro for helpful discussions.

## Supplementary Figures

Two distinct scene processing networks connecting vision and memory Christopher Baldassano, Andre Esteva, Diane M. Beck, Li Fei-Fei

**Supplementary Figure 1:**
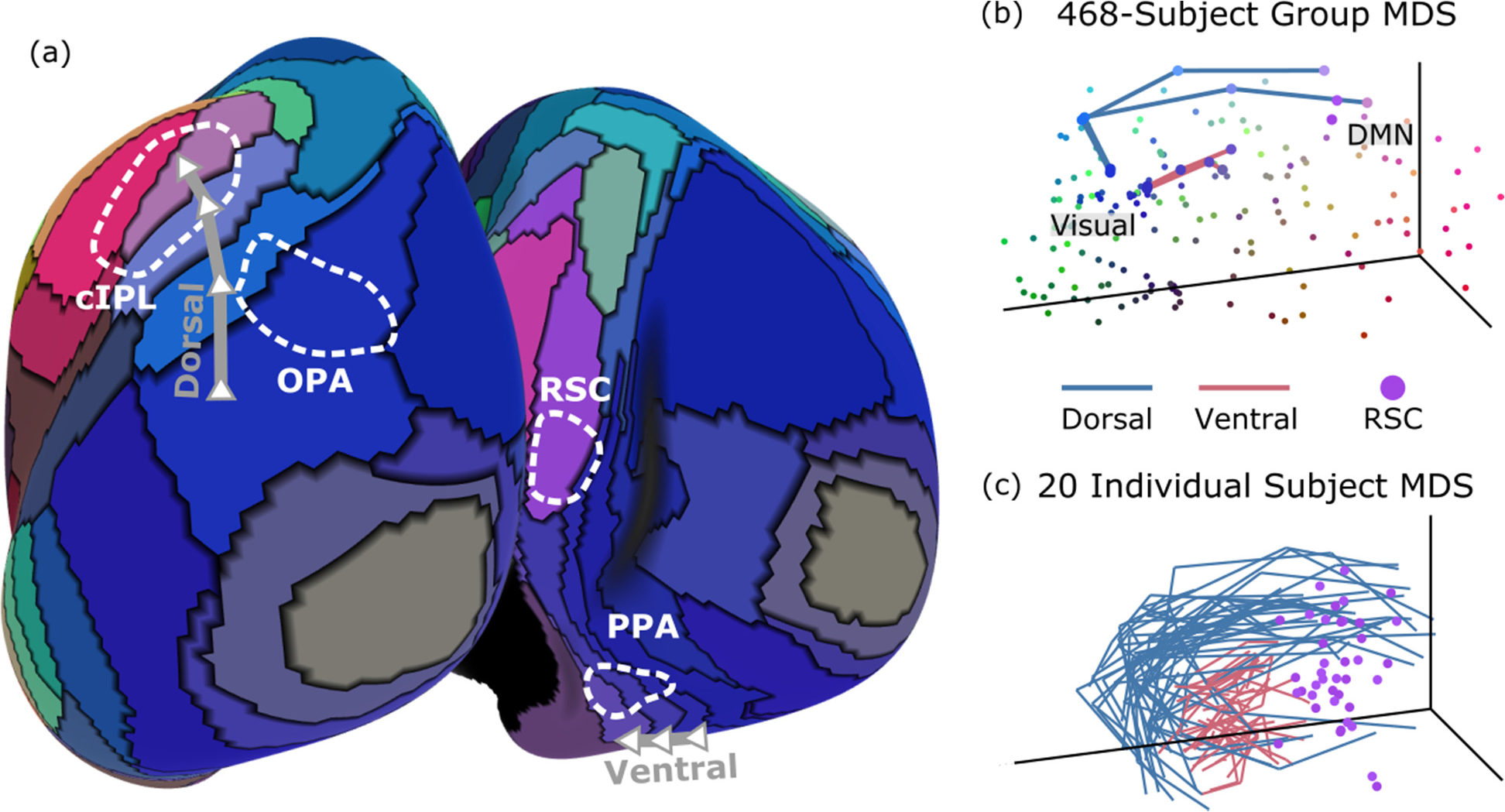
Multidimensional scaling of parcel connectivity matrix. We can use classical multidimensional scaling (MDS) to embed parcels into athreedimensional space RGB space. Distances in this space approximate the functional connectivity strength between parcels, such that strongly-connected parcels are close together in the embedding space and have similar colors. (a) There is a notable border in both dorsal (OPA to cIPL) and ventral (posterior to anterior PPA) cortex, where moving a short spatial distance along the cortical surface produces significant changes in functional connectivity properties. (b-c) These same paths are visualized in the MDS embedding space, both for the group and individual subjects. The most posterior regions show strong connectivity to other parcels in visual cortex, while the most anterior regions are instead more related to default mode regions.

**Supplementary Figure 2:**
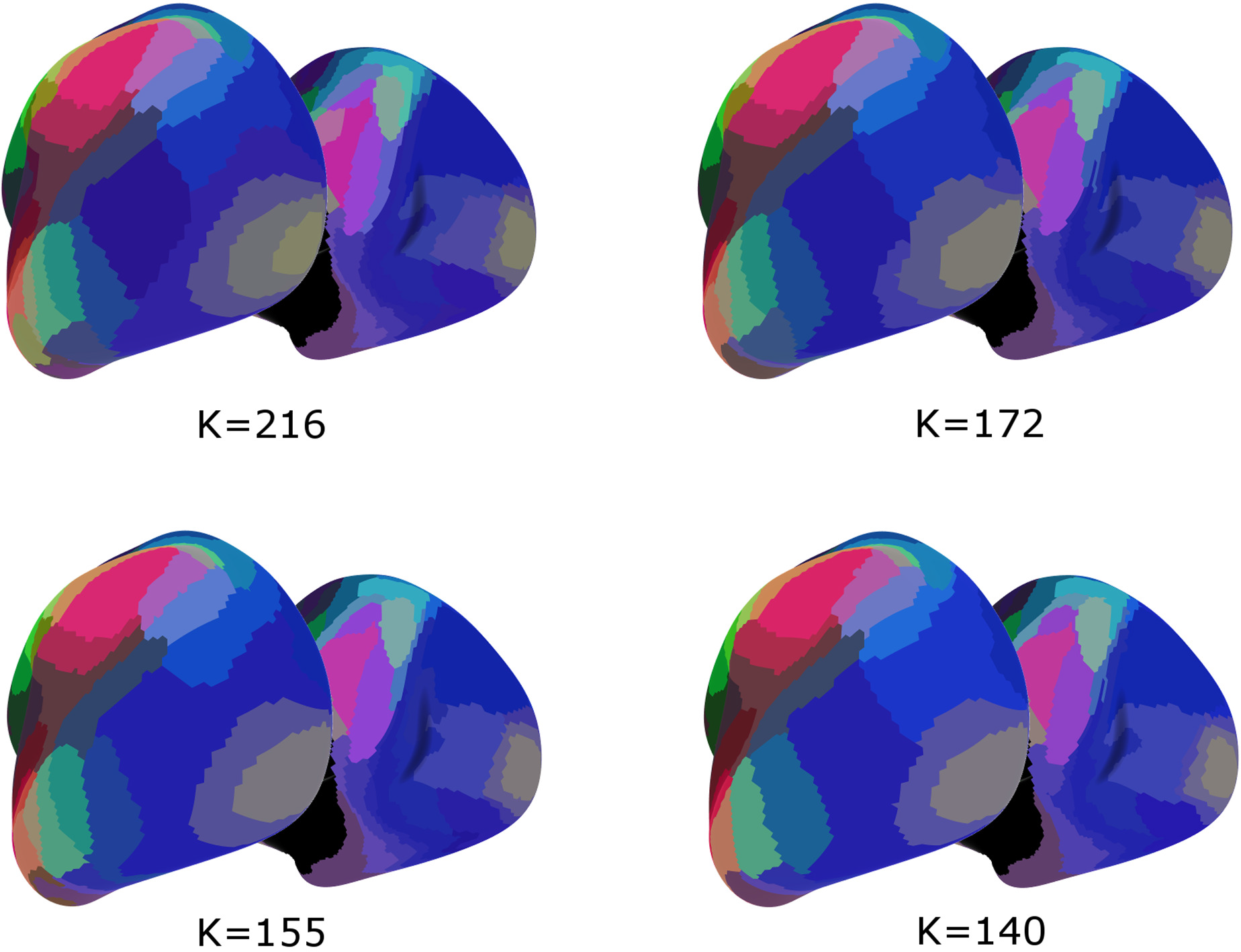
Varying the number of cortical parcels. The number of parcels chosen by our clustering algorithm (Baldassano et al., 2015) depends on a hyperparameter 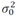 which controls the level of desired homogeneity within each cluster. Rather than using 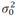=3000 as in the main text, we can reduce 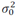 to 2000 (yielding 216 parcels) or increase 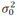 to 4000 (yielding 155 parcels) or 5000 (yielding 140 parcels). Performing the same MDS embedding as in Supplementary Figure 1 on each of these parcellations yields results that are qualitatively very similar, with the same transition between posterior and anterior scene regions.

**Supplementary Figure 3:**
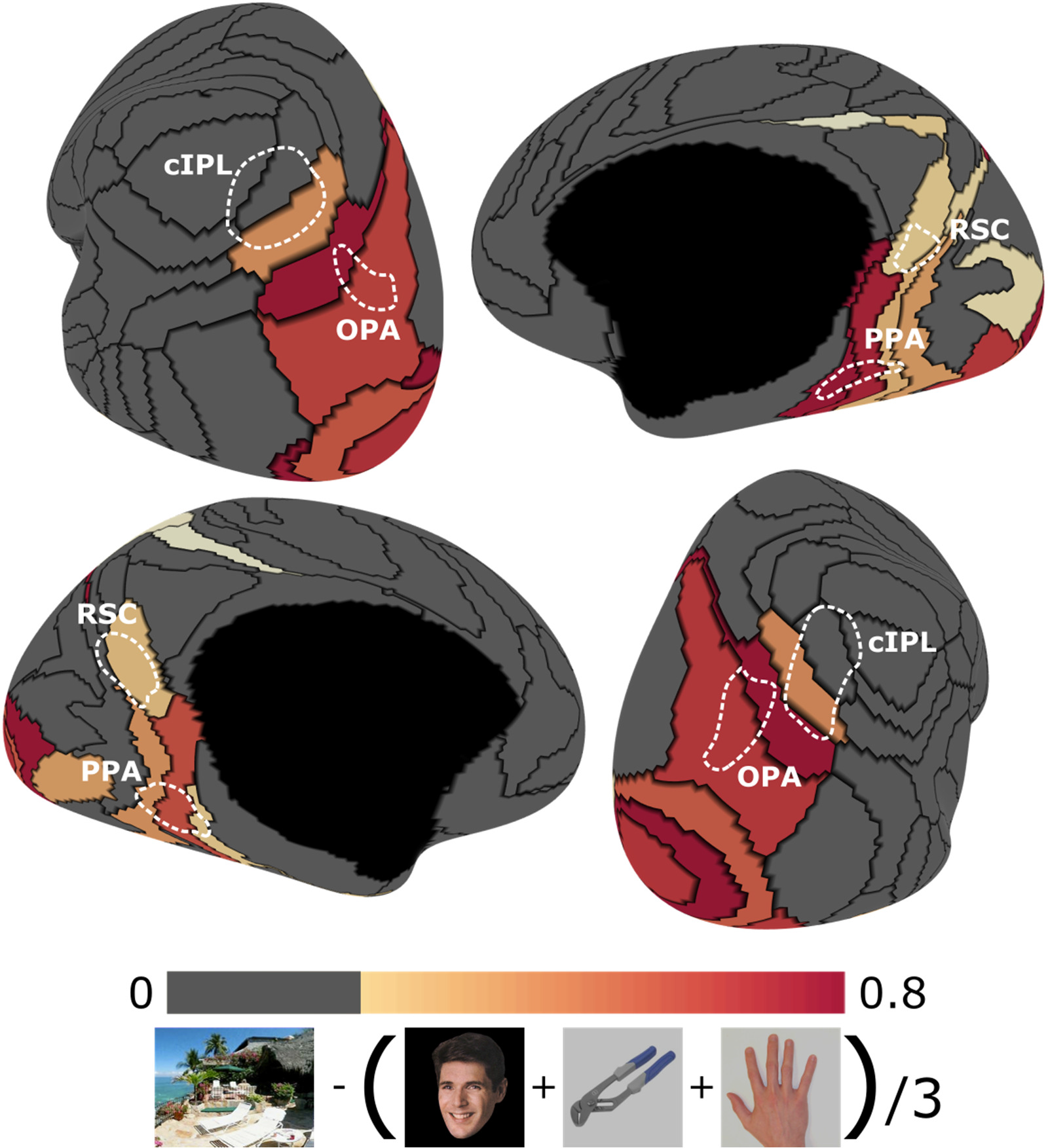
Testing for scene-sensitivity in HCP subjects. In the main text we use functional ROIs (OPA, PPA, RSC) defined using a standard localizer in an independent group of subjects. To validate that the parcels overlapping these ROIs are scene selective, we measured their meanresponseto scenes minus their response to other categories during the HCP working memory experiment. The results validate that these parcels are in fact scene selective in the HCP subjects. Note that early visual regions also show scene selectivity, likely because peripheral contrast was not well controlled in these stimuli. FDR<0.05.

